# AKT2 Loss Impairs BRAF-Mutant Melanoma Metastasis

**DOI:** 10.1101/2023.08.24.554685

**Authors:** Siobhan K. McRee, Abraham L. Bayer, Jodie Pietruska, Philip N. Tsichlis, Philip W. Hinds

**Affiliations:** Program in Genetics, Graduate School of Biomedical Sciences, Tufts University, Boston, MA 02111, USA; Department of Developmental, Molecular and Chemical Biology, Tufts University School of Medicine, Boston, MA 02111, USA; Program in Immunology, Graduate School of Biomedical Sciences, Tufts University, Boston MA 02111, USA; Department of Immunology, Tufts University School of Medicine, Boston, MA 02111, USA; Department of Cancer biology and Genetics, Ohio State University College of Medicine, Columbus, OH, 43210

**Keywords:** Melanoma, Metastasis, Cancer, AKT, Signaling

## Abstract

Despite recent advances in treatment, melanoma remains the deadliest form of skin cancer, due to its highly metastatic nature. Melanomas harboring oncogenic BRAF^V600E^ mutations combined with PTEN loss exhibit unrestrained PI3K/AKT signaling and increased invasiveness. However, the contribution of different AKT isoforms to melanoma initiation, progression, and metastasis has not been comprehensively explored, and questions remain whether individual isoforms play distinct or redundant roles in each step. We investigate the contribution of individual AKT isoforms to melanoma initiation using a novel mouse model of AKT isoform-specific loss in a murine melanoma model, and investigate tumor progression, maintenance, and metastasis among a panel of human metastatic melanoma cell lines using AKT-isoform specific knockdown studies. We elucidate that AKT2 is dispensable for primary tumor formation but promotes migration and invasion *in vitro* and metastatic seeding *in vivo*, while AKT1 is uniquely important for melanoma initiation and cell proliferation. We propose a mechanism whereby inhibition of AKT2 impairs glycolysis and reduces an EMT-related gene expression signature in PTEN-null BRAF-mutant human melanoma cells to limit metastatic spread. Our data suggest that elucidation of AKT2-specific functions in metastasis could inform therapeutic strategies to improve treatment options for melanoma patients.

**Graphical Abstract:** 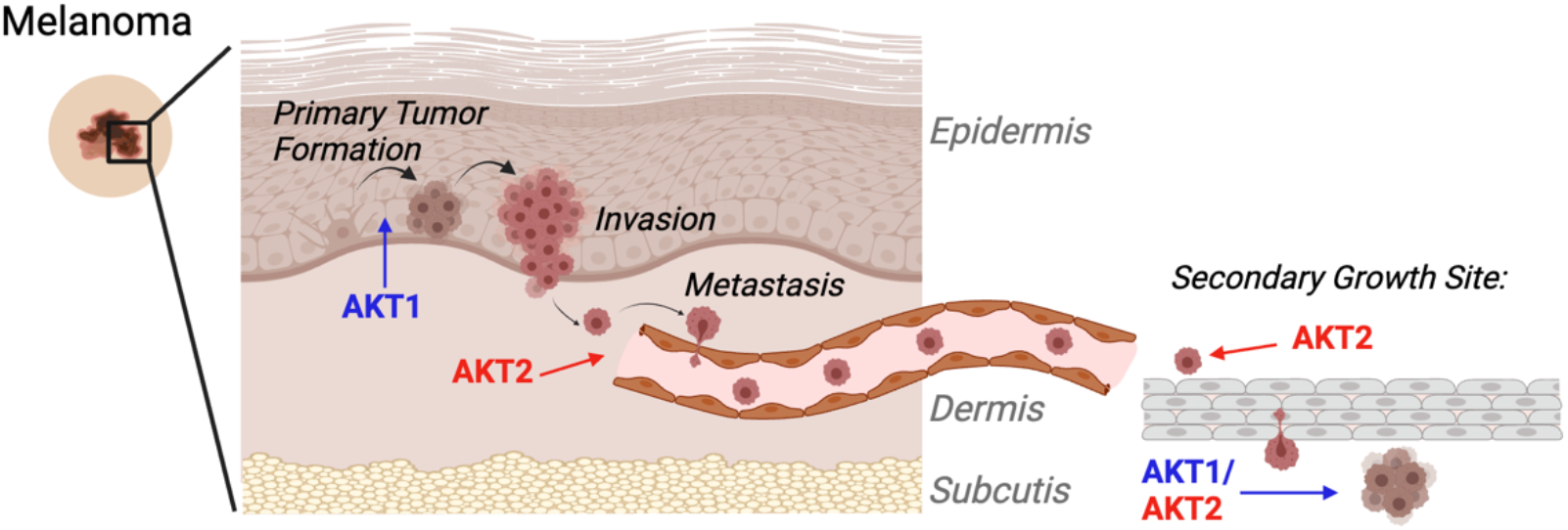

## 1. Introduction

Metastasis is responsible for the vast majority of all cancer deaths, yet no therapies preferentially target the metastatic process. A key hallmark of cutaneous melanoma is its rapid and efficient metastasis, facilitated in part by the PI3K/AKT pathway [1]. PI3K/AKT signaling is hyperactivated in late stage melanomas [2], often through loss of its negative regulator PTEN. PTEN inactivation leads to unrestrained activity of the PI3K effector kinase, AKT, which cooperates with pre-existing oncogenic BRAF mutations to promote metastasis [3]. Indeed, while the BRAF mutation V600E hyperactivates the MAPK signaling pathway and is a very common early mutation, it alone is insufficient for tumorigenesis and subsequent genetic and/or epigenetic alterations are required [4,5]. Alterations in PTEN, including inactivating mutations, epigenetic silencing, protein destabilization, or genetic loss, occur in as many as 50% of all melanomas, and correlate with advanced disease, including brain metastasis [6,7]. Accordingly, AKT is hyper-activated in melanoma brain metastases [8] and inhibition of the PI3K pathway effectively reduces metastasis in murine melanoma models [9]. Despite this, pan-PI3K inhibitors have yet to demonstrate clinical efficacy [10,11]. While six PI3K inhibitors have been approved for clinical use against breast cancer, hematologic cancers, and other diseases, only one exhibits pan-PI3K inhibition, while others are isoform specific. [12] A similar spectrum of clinical efficacy is likely from AKT inhibitors currently in clinical trials, with isoform specific inhibition better tolerated compared to pan-inhibition. This suggests that additional isoform specific PI3K/AKT inhibitors under development many be more effective therapeutically than pan-inhibition strategies. [13,14].

The AKT kinase family comprises three highly homologous yet functionally distinct isoforms (AKT1, AKT2, and AKT3) [15–18]. They differentially contribute to tumorigenesis in the breast, colon, and prostate [19,20] among other tissues, but isoform-specific inhibition has yet to be exploited as a therapeutic strategy. AKT isoforms achieve their specificity through differential activation, preferential substrate phosphorylation, and tissue distribution [19], but how these differential properties contribute to distinct cell transformation events is still being elucidated. While isoform specific effects of AKTs have been demonstrated in numerous other cancers [21–23] and in melanoma [24–27], many questions remain regarding the contribution of individual AKT isoforms to melanoma progression or metastasis, and few studies have interrogated the contribution of individual AKT isoforms to melanoma initiation. Large genomic datasets have provided some insights as to the relative frequency of AKT-isoform specific alterations; activating mutations or gene amplification of AKT1 and AKT2 are more common in BRAF^V600E^ mutant and PTEN-null melanomas [28,29] while AKT3 amplification is more common in BRAF^WT^PTEN^WT^ tumors [26], although AKT3 has also demonstrated cooperativity with BRAF^V600E^ to induce murine melanoma development [30]. With regard to melanoma metastasis, multiple and differing roles for AKT isoforms have been described in both mouse and human models. For example, upon loss of PTEN, invasiveness of human melanoma cell lines was enhanced through preferential phosphorylation of AKT2, while AKT3 phosphorylation reduced invasive potential of PTEN-null melanoma cells [31]. Similarly, it has been shown that the PHLPP1 phosphatase suppressed melanoma metastasis through dephosphorylation of AKT2 or AKT3, but not AKT1 in human melanoma cells [25]. In Braf^V600E^-driven murine melanomas lacking Cdkn2a, AKT1 activation was shown to enhance brain metastasis, while AKT2 and AKT3 had a less pronounced effect [27,32]. While the totality of data is suggestive that AKT isoforms play differing roles in melanoma initiation, progression, and metastasis, further clarifying studies are needed, especially given the importance of metastasis to melanoma outcomes.

Here we employ inducible AKT-isoform specific shRNAs and CRISPR/Cas9 mediated gene editing in human melanoma cell lines to outline a role for AKT2 in metastatic cell seeding of oncogenic BRAF-driven PTEN-null melanoma through enhancement of cell migration and invasiveness. As such, AKT2 depletion impairs and prevents metastasis, improving overall survival in mice. We further investigate AKT isoform phosphorylation in primary spontaneous murine melanomas and human metastatic melanoma cell lines, finding AKT2 phosphorylation specifically to increase in metastatic lesions. We further show that while AKT2 is dispensable for melanoma initiation, AKT1 contributes to BRAF-driven murine melanoma initiation and cell proliferation, in line with our previous results [33]. Mechanistically, we find AKT2 promotes metastasis and regulate glycolysis through a PDHK1-PDHE1α axis. This work supports continued investigation of targeted AKT inhibition as anti-melanoma therapy and presents a novel potential target in preventing metastatic spread.

## 2. Materials and Methods

### Mouse Strains

All mice were maintained in a heat- and humidity-controlled, AAALAC-accredited vivarium operating under a standard light-dark cycle. All protocols have been approved by the Institutional Animal Care and Use Committee (IACUC) at Tufts University School of Medicine, where mice were housed and experiments were conducted. BRAF^V600E^;Arf^-/-^ mice were bred in house, while AKT isoform knockout mice have been described previously [23]. NOD/SCID and C57Bl6/J mice were purchased from the Jackson Laboratory (Bar Harbor, ME).

### Xenografts

Male NOD/SCID mice (Jackson Laboratory, ME, 6-10 weeks old) were injected subcutaneously with 2x10^6^ human melanoma cells according to approved protocols. Once palpable, tumors were measured 3x weekly using calipers, and tumor volume was calculated using the formula [(π/6)*L*W^2^]. Tumors were allowed to grow until a limit of 1500mm^3^ or 2cm in any single direction was reached, or until mice became moribund. Doxycycline chow (200mg/kg, Teklad) was introduced when tumors were palpable.

### Luciferase Imaging

Mice were anaesthetized with isofluorane and injected intraperitoneally with 10 μl/g body weight of Luciferin (Fisher) 5 minutes prior to imaging. Imaging was performed using an IVIS SpectrumCT *in vivo* imaging system and analyzed using Living Image® Software.

### Metastasis Assays

Human (1x10^6^) or mouse melanoma cells (0.5x10^6^) were injected into the tail vein of NOD/SCID or C57Bl6/J mice, respectively. Mice were maintained on regular or doxycycline chow (200mg/kg, Teklad), imaged at 2 weeks post-injection and weekly thereafter. Mice were euthanized when moribund.

### Tumor Cell Isolation and Tissue Preparation

Tumors were minced and digested with 3 mg/ml collagenase and 250 U/ml hyaluronidase for 2-4 hours at 37°C. Contaminating red blood cells were lysed with RBC Lysis Buffer (Sigma), and organoids were triturated using an 18G syringe needle, incubated in 0.05% Trypsin/0.53mM EDTA, passed through a 40@m cell strainer, and cultured in RPMI1640/10% FBS/1% penicillin/streptomycin/fungizone.

### Cell Lines/Tissue Culture

SM1 cells were a generous gift of Antoni Ribas (UCLA) and TUMM cell lines were generated as described above. Human melanoma cell lines were generous gifts of Frank Haluska (Tufts Medical Center), and routinely validated for melanocytic identity by RNA or protein expression of MITF-M and pigment enzymes TYR and DCT, and tested for mycoplasma contamination. Human cells were maintained in DMEM (Invitrogen) with 10% FBS (Atlanta Biologicals) and 1% penicillin/streptomycin (Invitrogen). SM1 and TUMM murine cell lines were cultured in RPMI1640/10% FBS/1%penicillin/streptomycin/fungizone. Knockdown experiments utilized doxycycline (Sigma) at concentrations of 0.5-1 μg/mL. Doxycycline-inducible shAKT plasmids were generous gifts of Drs. Alex Toker and Rebecca Chin (Beth Israel Deaconess Medical Center). A non-targeting hairpin scramble sequence (Sigma) was cloned into the Tet-pLKO-puro backbone, a gift of Dmitri Wiederschain (Addgene plasmid #21915). 293T cells were transfected with shRNA constructs and packaging plasmids (psPAX2 and VSV-G) using PEI (polyethylenimine MW25,000, Polysciences). Viral supernatants were collected at 48h and 72h post-transfection and mixed. Stably transduced cell lines were generated by infection overnight in the presence of 8 μg/ml polybrene and 48 hours after infection were selected with 1 μg/mL Puromycin (Gibco) for three days. Cells infected with pLENTI-Luciferase-expressing virus (generous gift of Charlotte Kuperwasser, Tufts University) were selected with neomycin (G418, 500μg/mL, Gibco) for 2-3 weeks. CRISPR knockout WM1799 cells were previously generated and cultured as described.[33]

### Immunoblot and Immunoprecipitation Analysis

Cells were lysed in RIPA buffer containing protease and phosphatase inhibitors (Roche) and cleared by centrifugation. Protein concentration was determined by DC Protein Assay (Bio-Rad) and equivalent masses of protein were resolved using SDS-PAGE and transferred to 0.2 μm PVDF membranes (BioRad) for immunoblotting with indicated antibodies (see Table S3). Membranes were incubated with horseradish peroxidase-conjugated secondary antibodies and visualized using enhanced chemiluminescence (Pierce). For immunoprecipitation, samples were lysed in RIPA or CST lysis buffer (Cell Signaling Technologies), and protein concentrations were normalized to 1mg/ml and precleared with Protein A magnetic beads (Pierce). Antibody or equivalent control IgG was incubated overnight at 4°C, then with Protein A beads for 60-90 minutes. Antibody/bead complexes were washed extensively in lysis buffer, eluted by boiling in Laemmli buffer, and resolved by immunoblotting as described.

### Migration/Invasion Assays

Cells were pre-incubated for 24hr with Doxycycline (DOX, 1μg/mL) or DMSO for 24hrs, then seeded (25,000 cells) in the upper chamber of trans-wells (8 μm pores, Corning) in serum-free DMEM with DOX or DMSO, using 10% FBS as a chemoattractant and incubated overnight. Inserts were washed with PBS, scrubbed, fixed in ice-cold methanol, and stained with DAPI. Invasion assays utilized growth factor-reduced (GFR) Matrigel-coated transwells (Corning) and a 36 hr incubation.

### Anchorage Independent Growth (Soft Agar) Assay

Sterile low melting agarose (SeaPlaque, Lonza) was prepared at 5% stock concentration with 1% final concentration (bottom layer) prepared by diluting in appropriate cell culture medium, allowed to solidify at room temperature for 30 minutes, and overlaid with 10,000 cells per 6-well in 0.5% agarose. After solidifying 30 minutes, wells were overlaid with 0.5mL of media containing 0.5μg/mL of DMSO or Doxycycline and allowed to incubate at 37 degrees for 2-3 weeks, or until macroscopic colonies are visible. Media was refreshed every 2-3 days. Plates were then fixed with 10% neutral buffered formalin, washed 1x with PBS, and stained with 0.05% crystal violet overnight and washed with PBS until clear. Images were taken at 10x magnification and quantified using ImageJ.

### Seahorse Glycolytic Rate Assay

WM1799 cells were pre-incubated with DMSO/DOX for 48hrs, and 21,000 cells were seeded into microtiter plates (Agilent) one day prior to assay. One hour prior to assay, cells were washed and incubated with RPMI at pH 7.4 (Agilent) and placed in a non-CO2 incubator. Assays were performed using a Seahorse XFe96 Analyzer and Wave Software 2.4.0.

### Quantitative RT-PCR

Cells were lysed in TRIzol (Invitrogen) and RNA isolated by phenol-chloroform extraction according to manufacturer’s instructions, followed by lithium chloride/isopropanol precipitation. cDNA synthesis was performed using SMARTScribe™ Reverse Transcriptase (Takara, Inc) and qPCR was performed on a CFX96 real-time thermal cycler (Bio-Rad). See Supplementary Table 4 for primer sequences.

### Cell Cycle Analysis

Cells were cultured in DMSO or DOX as indicated, collected and washed with cold PBS, then fixed in ice-cold 70% EtOH for 30 min at 4 °C. Cell pellets were washed in staining buffer (PBS no Ca/Mg, 3% FBS, + mM EDTA), then incubated in staining buffer with 80 μg/mL Propidium Iodide and 0.125 mg/mL RNAse A (Thermofisher) for 40 min at 30 °C. Samples were run on a BD LSRII (BD Biosciences) and analyzed using FlowJo software.

### Statistical Analysis

Statistics were performed using GraphPad Prism 5.02, utilizing Student’s unpaired T-Test, One Way-ANOVA with Tukey post-test, or Kaplan Meyer Survival analysis as indicated. Significant p-values are listed, or noted as *P<0.05, **P<0.01, or ***P<0.001. Error bars represent standard error means.

## 3. Results

### AKT2 Depletion Impairs Cell Migration and Invasion in Human Melanoma Cells

It is well known that AKT phosphorylation, a surrogate marker for active AKT, increases with disease stage in clinical samples [34], but there is a paucity of information regarding the relative contributions of individual isoforms to disease progression. To investigate this, we first characterized AKT phosphorylation status across a panel of human melanoma cell lines to look for any preferential isoform activity at baseline *in vitro*. Immunoblotting for total phospho-AKT at both activating sites (Ser473 and Thr308), and isoform-specific phosphorylation (Summarized in supplementary Figure 1A-B) revealed a wide range of AKT phosphorylation levels, with no single AKT isoform preferentially phosphorylated (Supplementary Figure S1B). Further, as expected, there was an inverse correlation between PTEN expression and total AKT phosphorylation (Figure S1C).

**Figure 1.**
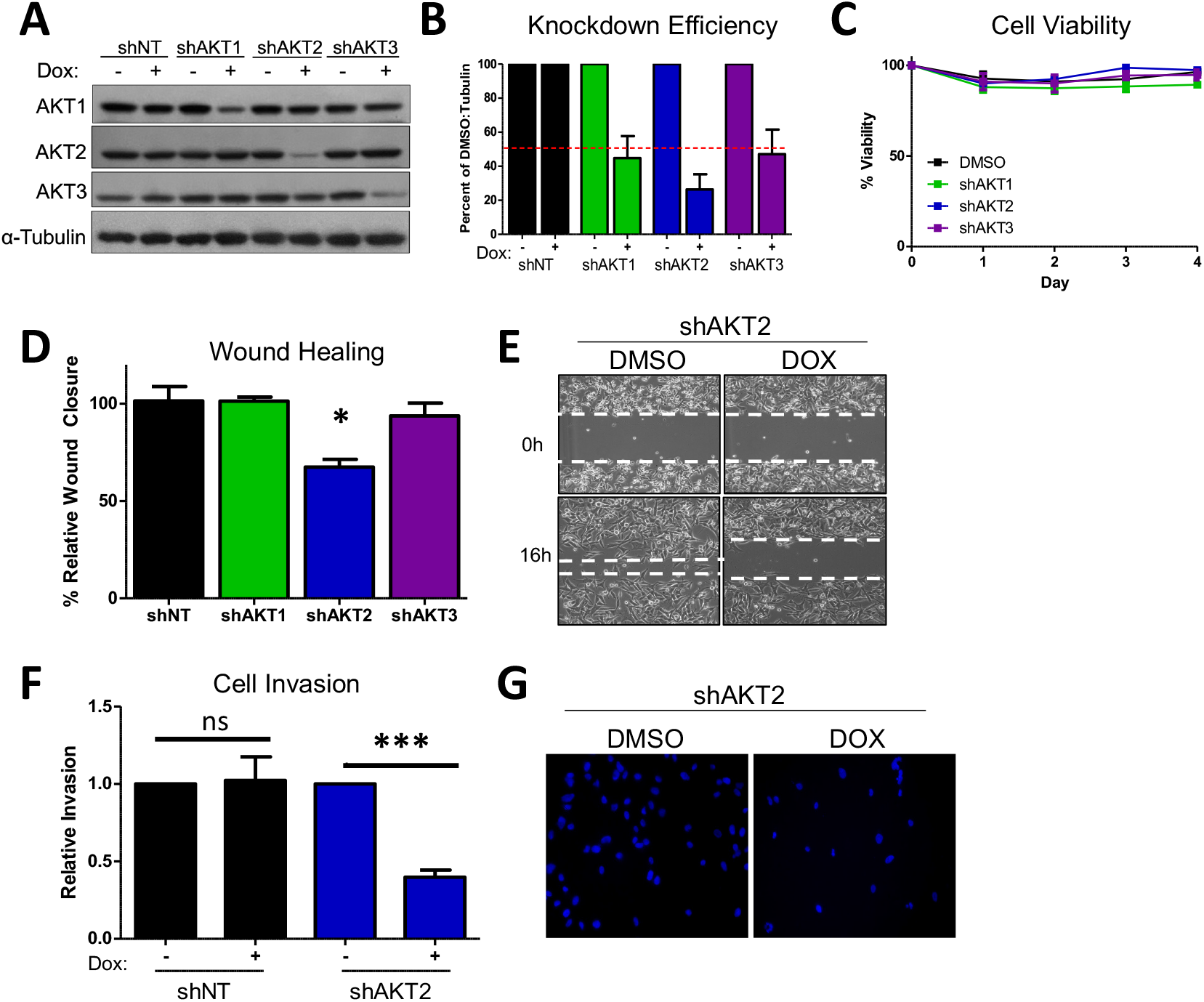
AKT2 depletion impairs migration and invasion in WM1799 human melanoma cell lines. A) A representative immunoblot of WM1799 human melanoma cells after stable cell line generation. Dox-inducible AKT-isoform knockdown (shAKT1/2/3) or non-targeting hairpin (shNT) expressing cell lines are shown after 72h treatment with 0.5ug/mL doxycycline. B) Quantitation of KD efficiency from three independent experiments using ImageJ to quantify intensity of bands normalized to percent of DMSO-treated total protein and loading control. C) Cell lines from (A) were grown in the presence of DOX or DMSO and collected at indicated times for cell counting with Trypan Blue exclusion to assess cell viability from n=2 independent experiments. D) Quantitation of wound closure in shWM1799 cells at 0h and 16h post wounding treated with DMSO-or DOX-containing media (4x magnification) and representative images (E) of shAKT2 WM1799 cells treated with DOX relative to DMSO vehicle treated cells. F) Quantitation of invasion ability of control (NT) or AKT2 KD (shAKT2) cells with or without DOX treatment as indicated. G. Representative images of DAPI-stained AKT2 KD cells on the underside of a Matrigel-coated membrane (20x magnification) following treatment with DMSO or DOX (left).

To investigate a possible contribution of AKT isoforms to metastatic potential in human cell lines, we generated luciferized doxycycline-inducible shRNA hairpins to AKT1, AKT2, and AKT3 [35] as well as a non-targeting hairpin (shNT), that efficiently reduced protein expression in the majority of melanoma cell lines from our panel, (Figure 1A-B, Figure S1D-S1F) without affecting viability (Figure 1C). We focused on cell lines in which all three AKTs exhibited detectable phosphorylation (WM1799, UACC903, and WM455), and then sought to test individual functions contributing to metastasis in these cells. Using a conventional *in vitro* wound healing assay, only AKT2 depletion but not AKT1 or AKT3 depletion, inhibited cell migration in three different human melanoma cell lines (Figure 1D-1E, Figure S2A-S2D). Next, we utilized a complementary trans-well assay, and observed AKT2 depletion reduced cell migration in response to a serum gradient in multiple cell lines (Figure S2E, S2G). Further, invasion through Matrigel was impaired by AKT2 depletion in the same human melanoma cell lines (Figure 1F-1G, Supplementary Figure S2F, S2H). This reduction was not due to defects in cellular proliferation, as AKT2 did not impair cell proliferation in any cell line as assessed by cell counting with trypan blue exclusion (Figure S5). Together, these data suggest that AKT2 depletion *in vitro* impairs functions required for melanoma metastasis such as migration and invasion.

### AKT2 Depletion Restricts Anchorage Independent Growth *In vitro* and *In vivo*

Anchorage-independent growth is required for the growth of metastatic cells; therefore, we interrogated the role of AKT2 in this process. Focusing our efforts on WM1799 cells, we seeded shAKT2 transduced WM1799 cells in soft agar overlaid with either DMSO-or doxycy-cline-containing media to knock down AKT2. We observed that AKT2 depletion reduced colony number (Figure 2A-2B) without affecting colony size (Figure 2C), indicating that AKT2 depletion limits the ability of cells to grow in 3D culture. Next, we assessed the effect of AKT2 depletion on the growth of WM1799 cells as subcutaneous tumors *in vivo*, which similarly requires anchorage independent growth in early tumorigenesis. We injected WM1799 shAKT2 cells subcutaneously into immunodeficient NOD/SCID mice (to exclude changes in melanoma cell growth due to an adaptive immune response) and allowed palpable tumors to form before transitioning a subset of these mice to doxycycline chow (Figure 2D). AKT2 knockdown significantly slowed tumor growth relative to control mice fed regular chow (Figure 2E), but tumors eventually grew in both groups, despite persistent AKT2 knockdown (Figure 2F) in the majority of tumors. Further, we observed no change in total AKT phosphorylation across tumors in which AKT2 was depleted, suggesting AKT1 or AKT3 activity may compensate over time to promote tumor growth.

**Figure 2.**
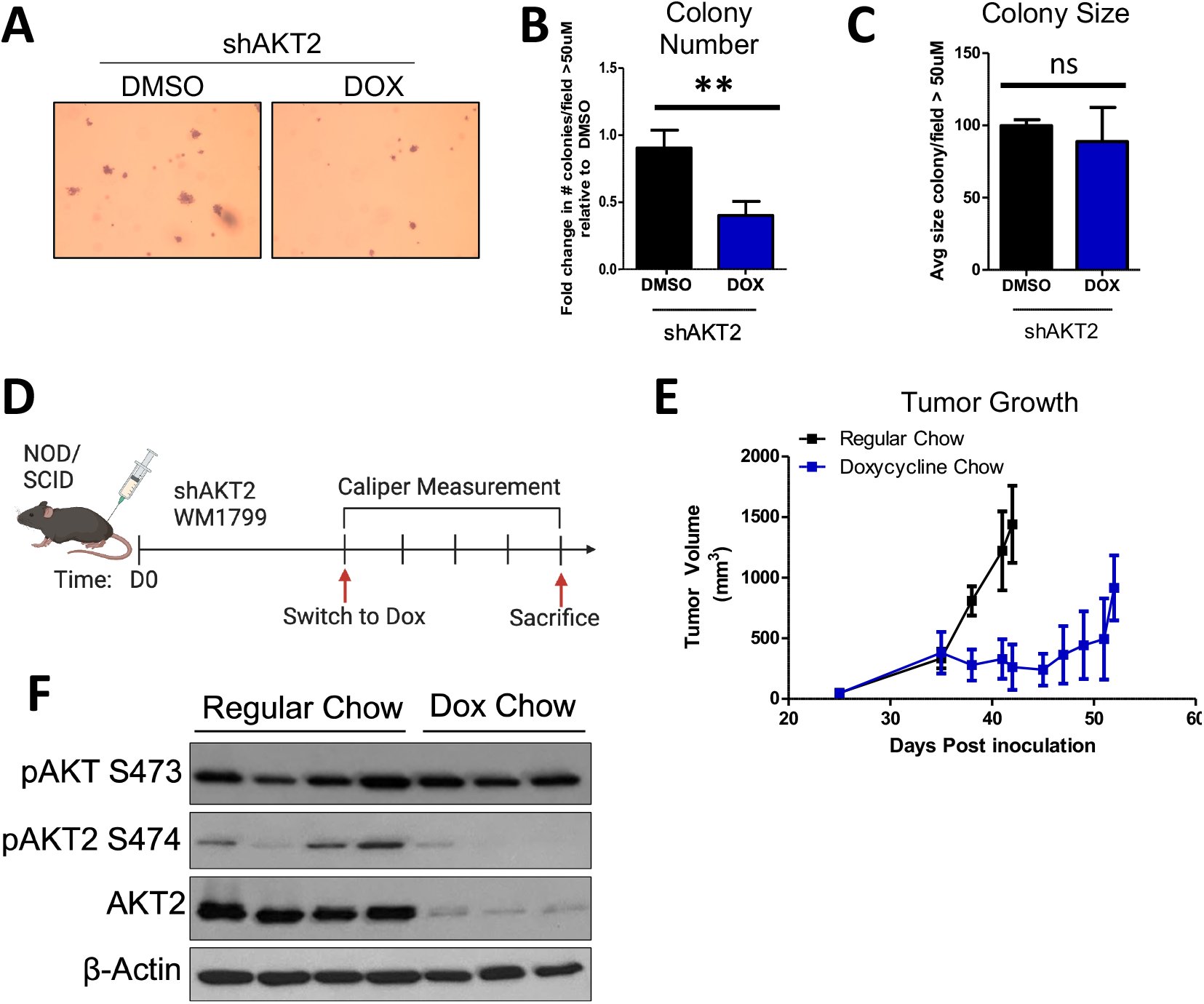
AKT2 depletion restricts anchorage independent growth. A) Anchorage-independent growth was assessed by seeding WM1799 AKT2 KD cells in soft agar and culturing in DMSO-or DOX-containing media for three weeks, after which time cells were fixed and stained with crystal violet (left, representative image at 4x magnification). B-C. Colonies greater than 50μM were counted and quantified using ImageJ. D. Schematic of experimental design to assess tumor growth potential. WM1799 shAKT2 cells were injected subcutaneously in NOD/SCID mice and allowed to form palpable tumors (500m3), then mice were randomized into groups receiving either Regular or DOX-containing chow to induce AKT2 KD (n=3-4 mice per group). E. Tumor volumes were determined using calipers at the indicated times, and mice were sacrificed when tumors reached 1500m3 in size. F. Tumors isolated from individual mice receiving either DMSO or DOX chow were subjected to immunoblotting for AKT2 protein and phosphorylation using beta-actin as a loading control.

### AKT2 depletion delays metastatic onset and extends survival of melanoma-bearing mice

To determine whether AKT2 is important for the process of metastatic seeding and metastatic nodule growth, we performed a tail vein metastasis assay in which WM1799 shAKT2-Luc melanoma cells were allowed to seed the lungs. This was followed by initiating AKT2 depletion 24 hours later by switching some mice to doxycycline-containing chow compared to mice fed regular chow (Figure 3A). By performing immunoblot analysis using lysate derived from isolated pulmonary metastases, we confirmed stable and robust AKT2 depletion occurred only in doxycycline-fed mice (Figure 3B). Using weekly imaging we found control mice fed regular chow displayed advanced metastatic disease at 6 weeks, with tumors observed at multiple sites, however mice fed doxycycline chow displayed only occasional tumor nodules at 6 weeks, albeit with significantly reduced size and frequency compared to control mice (Figure 3C-3D). While AKT2 depletion after metastatic seeding significantly improved overall survival in doxycycline chow fed mice versus regular chow mice, doxycycline-fed mice did succumb to eventual metastatic disease, suggesting that AKT2 reduction only delayed growth of metastatic lesions, and other AKT activity might compensate long term (Figure 3E).

**Figure 3.**
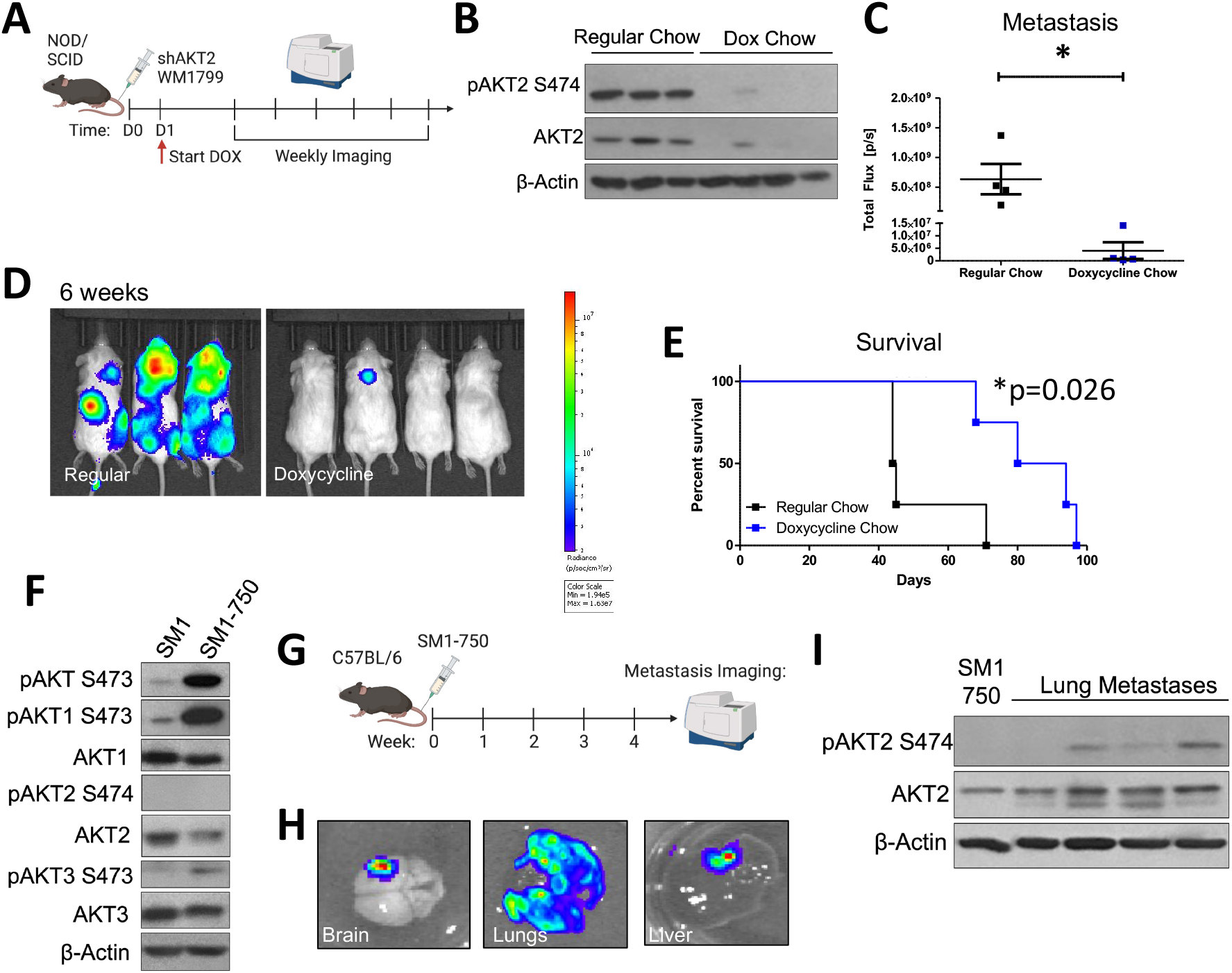
AKT2 depletion delays metastatic onset and extends overall survival of melanoma-bearing mice. A) Schematic representation of experimental design. Luciferized WM1799 cells expressing DOX-inducible shAKT2 hairpins were injected into the tail vein of NOD/SCID mice. Mice were started on DOX chow 1d post-injection and maintained on regular chow or DOX chow and monitored weekly IVIS SpectrumCT imaging. B. Immunoblot analysis of total and phospho-S474 AKT2 (P-AKT2) levels in metastatic tumors isolated from mice fed regular or doxycycline chow. C. Quantification of mice bearing metastases at 6 weeks post-injection and representative images (D) of luminescence 6 weeks after cell injection in mice fed doxycycline chow compared to regular chow (N=3-4 mice per group). E. Percent survival of mice bearing metastases F. SM1 and SM1-750 cell lines were established from spontaneous primary murine melanomas expressing BRAF^V600E^ and passaged in C57Bl6/J mice, then analyzed by immunoblotting. G. Schematic of experimental design of *in vivo* metastasis assays. SM1-750 cells expressing luciferase were injected into the tail vein of C57Bl6/J mice and tissues were analyzed using IVIS SpectrumCT imaging at 4 weeks post injection. H. Lungs, liver, and brains collected from mice were imaged ex-vivo, and representative images from n=4 mice are shown. I. Lung metastases were excised from 4 individual mice (at least 3-4 nodules per mouse were combined) and homogenized to generate protein lysates and analyzed by immunoblotting.

### AKT2 Phosphorylation Occurs in Metastatic Mouse Melanoma Lesions

To study AKT2 in the context of metastatic murine melanoma, we generated an aggressive murine melanoma cell line, SM1-750, which exhibits high metastatic potential. The SM1 cell line was previously derived from melanomas arising in a BRAFV600E;Arf-/-mouse and is tumorigenic in syngeneic mice [36]. In our hands, subcutaneous tumor formation by SM1 cells in the C57BL6/J strain occurred in only a small subset of injected mice. To increase the tumor “take rate,” we passaged the SM1 cell line in tumor-forming C57BL6/J mice through several rounds of injection and tumor formation, resulting in isolation of the SM1-750 line, which displays a nearly 100% take rate. Subsequently, the SM1-750 line was engineered to express luciferase, and maintenance of the ability of these cells to form tumors was confirmed. (Figure 3F, Figure S3).

SM1 cells exhibit significant AKT1 and AKT3 phosphorylation but nearly undetectable AKT2 phosphorylation, and the newly derived SM1-750 cells showed further elevated phosphorylation of only AKT1 and AKT3. SM1-750-cells were injected into the tail vein of syngeneic mice and the injected mice were imaged after luciferin injection to visualize metastatic progression (Figure 3G). Mice readily developed metastatic lesions, which could be visualized in the lungs, brain, and liver (Figure 3H). Spontaneous brain metastases are relatively rare in murine melanomas, and have previously been linked to AKT1 activation [27]. Interestingly, immunoblotting of AKT isoform phosphorylation in discrete lung tumor metastases from individual mice revealed that AKT2 activation (indicated by S474 phosphorylation) is dramatically increased compared to that of primary tumors (Figure 3I). We further found AKT2 upregulation in metastatic lesions was not due to adaptive immunity-dependent selection as we also observed this in SM1-750 metastases isolated from immune-deficient Rag2^-/-^ mice (Figure S3A).

We then sought to characterize AKT isoform phosphorylation in spontaneously arising primary melanomas relative to murine melanomas with metastatic potential. We previously developed a mouse model of BRAF^V600E^-driven spontaneous melanoma in which melanocyte-targeted human BRAF^V600E^ cooperates with tumor suppressor p19^ARF^ loss (hereafter referred to as Arf^-/-^), to facilitate melanoma formation and in which AKT phosphorylation was observed in tumors but not normal skin [37]. The Arf tumor suppressor is commonly lost in human melanomas and melanoma penetrance in our model is increased on an Arf^-/-^ background [38]. We isolated cell lines from spontaneously arising primary tumors in these mice and investigated isoform-specific phosphorylation patterns (Tufts University Mouse Melanoma, TUMM, see Supplementary Figure 3B-3C). This analysis revealed ubiquitous total AKT phosphorylation on the activating residue serine 473 in TUMM cell lines as expected, as well as readily detectable AKT1 phosphorylation, with rare AKT3 phosphorylation and almost no AKT2 phosphorylation, in line with our observations in SM1-750 cells (Figure S3C). Further, neither TUMM nor SM1-750 cells showed AKT2 phosphorylation in low attachment cell culture plates or in primary tumors produced following subcutaneous injection into NOD/SCID mice (Figure S3D-S3E), indicating that while AKT1 or AKT3 phosphorylation can drive melanoma cell growth and primary tumor formation, activation of AKT2 correlates strongly with an ability to grow in the metastatic niche.

### Prophylactic AKT2 depletion prevents metastatic cell seeding

To determine if a role for AKT2 in cellular migration and invasion as well as metastatic progression further extended to initial metastatic seeding, we pre-treated WM1799 shAKT2 cells with doxycycline-containing media compared to DMSO-containing media alone for 72 hours prior to tail vein injection into NOD/SCID mice, then maintained the mice on doxycycline containing food or regular chow for 6 weeks. Metastatic progression was then monitored by luminescence (Figure 4A). Mice maintained on regular chow displayed luminescent tumor nodules within 3 weeks, which progressed to advanced metastatic disease by 6 weeks, including multiple distant metastases under the forelimbs, the head, neck, and mesenteric lining (determined at autopsy, not shown), consistent with lymphatic dissemination (Figure 4B-C). Contrary to control mice, mice that received shAKT2 cells and doxycycline chow remained healthy, with no detectable tumors by 6 weeks (Figure 4B-C). Additionally, when maintained on doxycycline chow, these mice were protected from detectable metastasis, even up to 12 weeks (Figure 4D-E). These findings suggest that AKT2 is either required for metastatic seeding or growth of seeded cells but does not distinguish whether AKT2 KD cells were eliminated from circulation or were simply latent in mice fed doxycycline chow. To address this question, we asked if tumors would emerge after removal of doxycycline chow. A subset of mice injected with WM1799 shAKT2 Luc cells and fed doxycycline chow were switched to regular chow after the first 6 weeks, at which time they still did not have detectable metastases. Mice were monitored weekly, and after an additional 6 weeks, none of the mice removed from doxycycline chow developed metastases (Figure 4F-G). These results suggest that prophylactic targeting of AKT2 impairs seeding of invasive cells in the metastatic niche, fully preventing metastatic formation rather than restraining growth of dormant but intact metastatic tumor cells.

**Figure 4.**
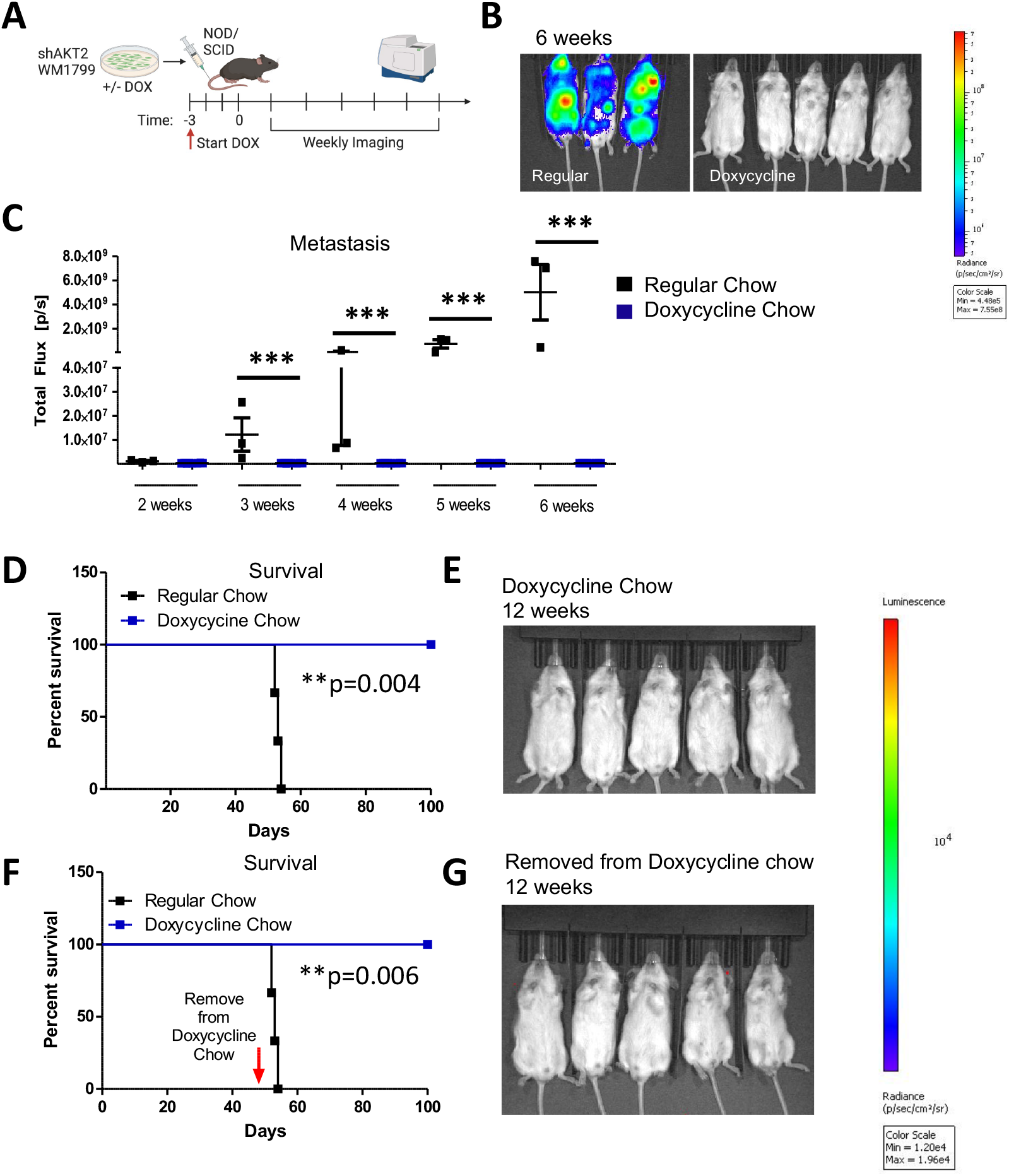
Prophylactic AKT2 depletion prevents metastatic cell seeding. A. Schematic representation of experimental design. AKT2-depleted (3 days of DOX treatment) or control (DMSO treated) WM1799 shAKT2 Luc cells were injected into the tail vein of NOD/SCID mice fed DOX chow or regular chow respectively for three days prior to injection (N=3-5 mice per group). Mice were provided regular or DOX chow and monitored weekly by IVIS SpectrumCT imaging. B. Image of mice at 6 weeks post-injection in DOX treated vs. regular chow mice. C. Quantification of luminescence from mice in (B) using LivingImage software at indicated times. D. Survival of regular-chow-fed (control) and DOX-fed mice injected with DMSO-or DOX-treated shAKT2 Luc cells, respectively. Mice were sacrificed when mori-bund. E) Luminescence assayed at 12 weeks in mice injected with AKT2 KD cells and fed Dox chow. F-G. A subset of mice treated as in D and E were switched to regular chow after 6 weeks on DOX chow and were assessed for the presence of metastases by IVIS SpectrumCT imaging after an additional 6 weeks and show no evidence of progressing metastatic lesions.

### AKT2 Deletion Impairs Melanoma Migration, Invasion, and Metastasis

To confirm these findings with a more robust depletion of AKT2, we utilized isoform specific CRISPR/Cas9 knockout in WM1799 cells in which we previously confirmed stable isoform specific knockout [33]. We first characterized the properties of AKT2 knockout (KO) WM1799 cells compared to non-targeting (NT) cells *in vitro* as done with the inducible knockdown cell lines (Figures 1-2). As expected and in line with our results using inducible knockdown, we found that AKT2 KO cells had impaired wound healing by scratch assay, migration through transwells, and Matrigel invasion, compared to NT cells (Figure 5A-5C), all indicative of impaired metastatic properties. To test this we further engineered NT and AKT2 cells to express luciferase and injected them into the tail veins of NOD SCID mice (Figure 5D). At 8 weeks postinoculation, NOD SCID mice that received AKT2 KO cells had significantly lower metastatic burden than those injected with NT cells (Figure 5E-5F). Further, mice injected with AKT2 KO cells showed a survival benefit relative to NT-injected mice (Figure 5G), consistent with previous results indicating that AKT2 depletion delays the onset of metastatic disease and overall survival.

**Figure 5.**
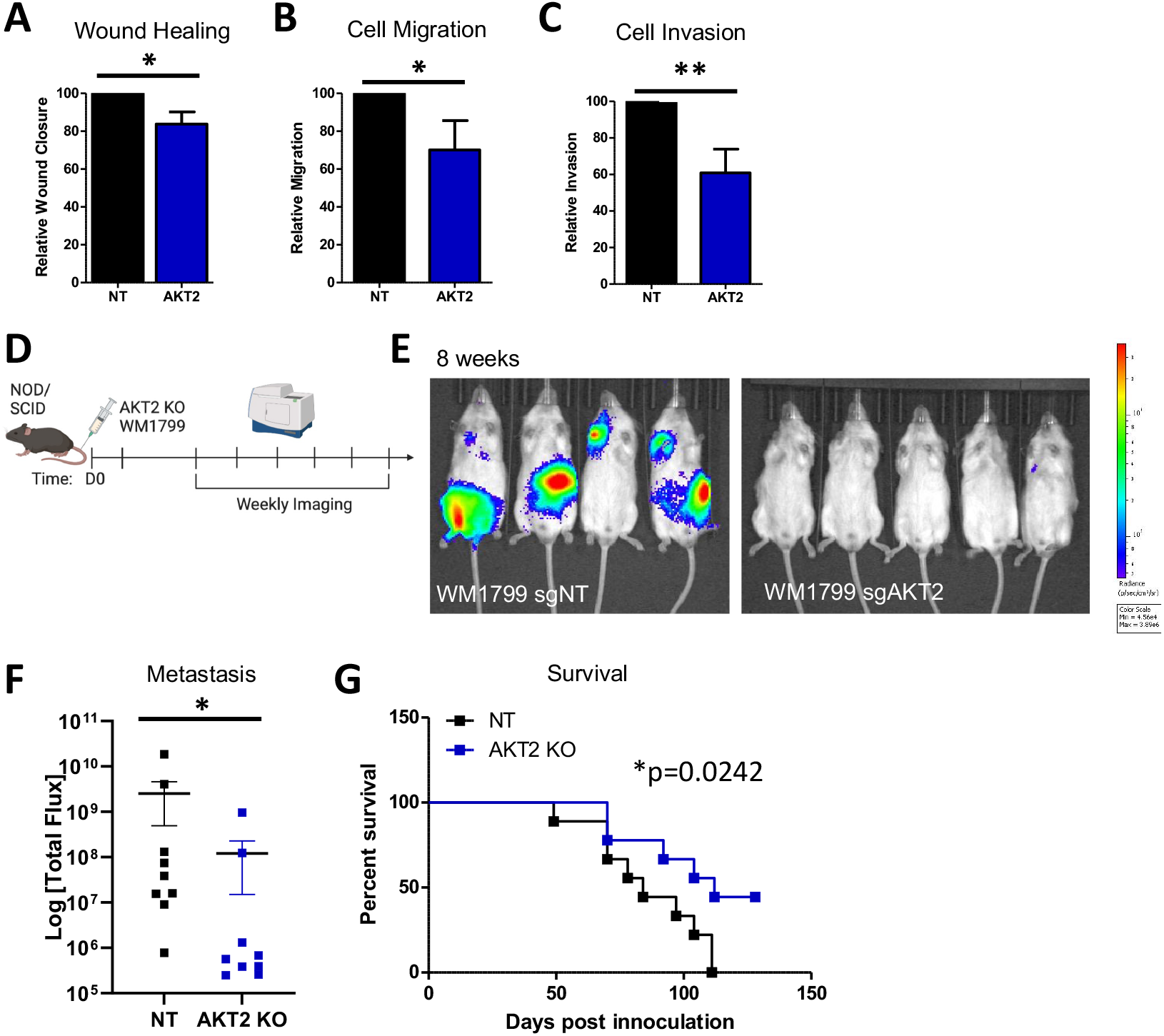
AKT2 Knockout impairs human melanoma cell migration, invasion, and metastasis. A-C. Ability of WT or AKT2 KO WM1799 cells in wound closure using scratch assay (A), cell migration using transwell assay (B), or cell invasion through a Matrigel coated membrane (C). D. Experimental schematic in which AKT2 KO WM1799 cells were engineered to express luciferase and injected into the tail veins of NOD/SCID mice, metastasis was monitored using IVIS SpectrumCT imaging, shown are representative mice at 8 weeks (E) with quantification (F). G. Overall percent survival of mice injected with WM1799 AKT2 KO cells compared to NT cells.

### AKT1 Deletion Impairs Primary Tumor Formation

We next sought to test if AKT2 knockout impaired tumor initiation compared to AKT1 and AKT3 deletion, as our CRISPR-edited WM1799 cells exhibited more robust AKT2 deletion compared to the inducible knockdown lines, which did not impact tumor initiation. We injected each isoform specific knockout into the flanks of NOD/SCID mice and monitored tumor formation and growth using caliper measurement (Figure S4A). Interestingly, we found that while AKT2 and AKT3 KO WM1799 cells developed tumors similarly to NT cells, only AKT1 KO cells had significantly delayed tumor growth. (Figure S4B).

To further investigate the impact of genetic loss of each AKT isoform on melanoma progression, we crossed melanoma prone BRAF^V600E^; Arf^-/-^ mice with AKT isoform knockout mice [23] to generate BRAF^V600E^; Arf^-/-^; AKT^-/-^ compound mutant mice (Breeding scheme, Figure S4C). As BRAF^V600E^; Arf^-/-^ mice have significant tumor formation and overall reduced survival [38], we analyzed the contribution of each AKT to long term survival. Again, AKT1 loss in the context of oncogenic BRAF and Arf loss provided a survival benefit for these melanoma prone mice, which was not observed with AKT2 or AKT3 loss (Figure S4D). This is in line with our data showing only strong AKT1 phosphorylation in primary mouse tumors, distinguishing the role for AKT2 in metastatic seeding from the role for AKT1 in primary murine melanoma formation.

### AKT1 Knockdown Impairs Cellular Proliferation and Anchorage Independent Growth

To investigate why AKT1 but not AKT2 impaired primary tumor growth, we analyzed cellular proliferation in our inducible AKT1 and AKT2 knockdown melanoma cell lines. We found that in WM1799, UACC903, and WM1158 cells, only AKT1 but not AKT2 knockdown impaired cellular proliferation after 4 days in culture with Doxycycline containing media (Figure S5A). We went on to perform cell cycle analysis in shWM1799 cells and discovered that Doxycycline treated AKT1 knockdown cells exhibited increased G1 arrested cells and fewer G2/M phase cells, while NT and AKT2 knockdown cells showed no cell cycle defects in the presence of Doxycycline (Figure S5B-S5C). AKT1 knockdown cells further showed significantly reduced BrdU incorporation in the presence of Doxycycline compared to NT and AKT2 cells (Figure S5D), confirming a role for AKT1 in cellular proliferation in line with existing literature [17], and one mechanism of delayed primary tumor formation.

We further tested whether AKT1 played a role in anchorage independent growth, as AKT1 KO WM1799 cells not only showed delayed tumor formation but also a reduction in tumor growth. Indeed, knockdown of AKT1 reduced colony formation in soft agar with a trend in decreased colony size (Figure S6A-S6C). We went on to test this observation *in vivo* by implanting shAKT1 WM1799 cells in NOD/SCID mice and switching mice to Doxycycline containing chow compared to regular chow (Figure S6D). Mice on Doxycycline chow had significantly reduced tumor growth compared to regular chow fed mice, together showing that AKT1 drives tumor cell proliferation while AKT1 and AKT2 both contribute to anchorage independent growth.

### AKT2 depletion inhibits EMT and impairs glycolysis through PDHK1 phosphorylation in melanoma cells

In order to investigate possible mechanisms whereby AKT2 could support metastatic growth and survival, we first interrogated the impact of AKT2 depletion on the epithelial-mesenchymal transition (EMT), an early step in the acquisition of invasive and metastatic capability[39]. Only AKT2 knockdown but not AKT1 or AKT3 knockdown in WM1799 cells reduced expression of pro-metastatic transcription factors ZEB1 and Snail, and the matrix-remodeling enzyme MMP2, concomitant with a trend in increased E-cadherin expression (Figure 6A). Additionally, only AKT2 knockdown decreased expression of TEA domain (TEAD) genes (Figure 6A), invasion-associated genes previously implicated in melanoma metastasis [40]. These data are consistent with an isoform-specific role for AKT2 in programming melanoma cells for EMT transition and subsequent metastasis.

**Figure 6.**
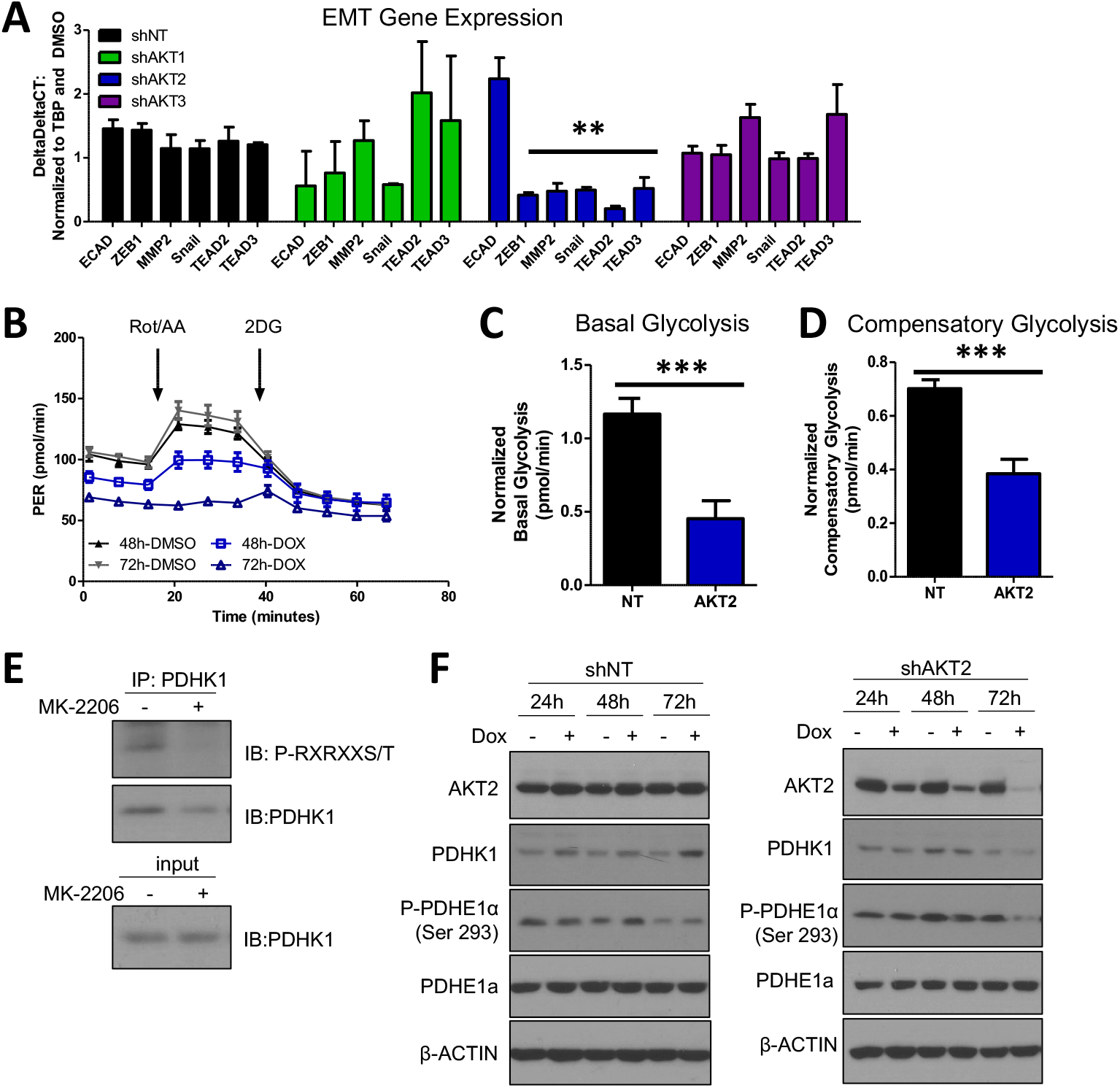
AKT2 depletion alters EMT and impairs glycolysis in melanoma cells. A. RT-qPCR analysis of EMT and invasion-associate transcripts in WM1799 shNT, AKT1 KD, AKT2 KD, or AKT3 KD cells after 48h of DOX treatment. Expression levels were normalized to shNT cells. B. Representative GRA plot shows proton efflux rate (PER) of WM1799 shAKT2 cells cultured in DMSO-or DOX-containing media for 48 or 72 hours, which was used to quantify basal glycolysis (C) and compensatory glycolysis (D) in DOX-treated WM1799 shNT cells. E. PDHK1 immunoprecipitation was performed after 24 hours of MK-2206 treatment, then immunoblot was performed for AKT consensus site phosphorylation, input immunoblot for PDHK1 shown in lower panel. F. Immunoblot analysis of PDHK1 expression or PDHE1a expression and phosphorylation in WM1799 shNT or AKT2 KD after treatment with DOX for indicated time points.

It has long been appreciated that tumor cells undergo metabolic reprogramming during tumor progression and especially during metastasis, with an increased reliance on glycolysis over oxidative phosphorylation, a phenomenon known as the Warburg effect [41]. To determine whether AKT2 knockdown directly disrupts glycolysis, we employed a Seahorse glycolytic rate assay (GRA), which quantitatively measures proton efflux rate (PER) in real time under standard conditions disruptive to mitochondrial metabolism. Using shNT or shAKT2 WM1799 cells, we performed the glycolytic rate assay after 48 or 72 hours of incubation with DMSO-or DOX-containing media (Figure 6B). Comparison of basal glycolytic metabolism in shNT and shAKT2 cells revealed that AKT2 knockdown suppressed basal glycolytic metabolism (Figure 6B-6C). 72 hours of AKT2 knockdown also suppressed glycolysis independent of mitochondrial respiration since Rot/AA injection was unable to increase PER (Figure 6B). Compensatory glycolysis was also significantly reduced in AKT2 knockdown cells compared to DOX-treated WM1799 shNT cells (Figure 6C). Together, these results strongly suggest that selective AKT2 knockdown inhibits glycolysis in WM1799 human melanoma cells.

To explore the potential mechanisms underlying the observed AKT2-dependent glycolytic defect, we evaluated the impact of AKT2 suppression on key enzymatic regulators of glycolysis. Pyruvate dehydrogenase kinases (PDKs) are critical regulators of the pyruvate dehydrogenase complex (PDC) through inactivating phosphorylation events, with subsequent decreases in TCA cycle activity shifting towards anaerobic respiration [42]. Of the four PDK isoforms (PDHK1-4), PDHK1 is the most frequently studied in the context of tumorigenesis, and has previously been shown to be a direct target of AKT2 in prostate adenocarcinoma cells [42]. We first confirmed that PDHK1 was an AKT target in WM1799 cells by performing immunoprecipitation of PDHK1 in the presence or absence of the AKT inhibitor MK2206 and probing for phosphorylation of the AKT consensus site, showing AKT specific phosphorylation on isolated PDHK1 that was eliminated by MK2206 (Figure 6E).

We further tested the effect of AKT2 knockdown on PDHK1 levels and found shAKT2 cells treated with Doxycycline resulted in decreased levels of PDHK1 by 72 hours that was absent in WM1799 shNT cells (Figure 6F). We then tested the effect of this on the PDHK1 target pyruvate dehydrogenase E1 component subunit alpha (PDEH1α). We found that the drop in PDHK1 at 72 hours of Doxycycline treatment in shAKT2 cells resulted in a loss of PDEH1α phosphorylation, which would be expected to shift metabolic flux away from glycolysis back to the TCA cycle, one mechanistic explanation for the loss of glycolytic activity in shAKT2 cells. Overall, our findings are consistent with a role for AKT2 in the regulation of not just EMT, but also glycolytic vs. aerobic respiration, promoting melanoma invasiveness and metastatic activity.

## 4. Discussion

Despite clear evidence that the PI3K/AKT pathway is important for tumor progression and metastasis in melanomas, effective therapeutic inhibition of the AKT pathway has been challenging [10]. A potential pitfall in targeting the AKT pathway may be the strategy of pan-inhibition despite strong evidence of AKT-isoform-specificity in cancer. This study sought to investigate the contribution of AKT isoforms to melanoma initiation and metastasis, utilizing both murine and human melanoma models.

Although it is well-known that AKT activation drives tumor progression and metastasis in melanoma [43–46], the contributing AKT isoform(s) were not determined. Initially, we crossed mice with BRAF-driven murine melanoma previously developed in our lab [38], to mice lacking different AKT isoforms [23] and monitored the mice derived from these crosses for melanoma development. We observed that AKT1 loss extended overall survival in melanoma prone mice, while AKT2 and AKT3 loss did not (Figure 1C). This suggests that AKT1 promotes melanoma initiation and growth, consistent with a well-described role for AKT1 in tumor promotion [15,21] and cell proliferation and survival [17]. This is also consistent with our observation that tumor-derived cell lines consistently show robust phosphorylation of AKT1 compared with other AKT isoforms (Figure 1A, D), suggesting AKT1 but not AKT2 or AKT3 plays a critical role in melanoma initiation. One caveat to these findings is that 20% of BRAF^V600E^/Arf-/-mice die for reasons unrelated to melanoma [38], however further studies would need to be done to determine exact cause of death. As such, melanoma-free survival, or exclusion of animals that both lacked melanoma and also died for reasons unrelated to melanoma would more accurately reflect the relative contributions of AKT isoforms to melanoma initiation and tumor promotion. However, because early primary melanomas pose a relatively low clinical risk, we focused our continuing studies on metastatic disease progression, which still presents the greatest treatment challenge for patients.

To address the potential involvement of AKT isoforms in metastasis, we chose to examine how the loss of AKT isoforms effects the metastatic potential of a melanoma cell line we established from a BRAF^V600E^/Arf^-/-^ mouse melanoma. The cell line was engineered to express luciferase and following IV inoculation in syngeneic immunocompetent mice, it could be monitored by bioluminescence. Using this approach, we discovered that increased phosphorylation of AKT2 was present in metastatic lesions, despite the lack of such phosphorylation in primary tumors and parental cell lines, suggesting that AKT2 activation may facilitate metastatic seeding, survival and/or growth of the cells seeded in metastatic sites. We have previously reported a low but significant rate of spontaneous lung metastasis in the BRAF^V600E^/Arf^-/-^ melanoma model [38] Based on this observation, future studies could use this model to determine whether the AKT isoforms differentially control the rate of metastatic melanomas. This could be addressed by monitoring the metastatic burden of BRAF^V600E^/Arf^-/-^/AKT^-/-^ mice lacking individual AKT isoforms.

The preceding data suggest that AKT2 differentially promotes the metastatic potential of melanomas, but in order to address the mechanism, we utilized a set of human melanoma cell lines driven by mutation in BRAF, as well as by PTEN loss. Functional loss of the tumor suppressor PTEN leading to AKT activation occurs in a large proportion of melanomas [7] and frequently co-occurs with oncogenic BRAF mutations [26]. However, the isoform-specificity of AKT in the context of PTEN loss has remained largely unexplored. Therefore, using several BRAF^V600E^ metastatic human melanoma cell lines with PTEN loss in which all three AKT isoforms are phosphorylated (Supplemental Figure 2A-B), we investigated the consequence of isoform-specific knockdown on cell migration and invasion. Our results show that AKT2 depletion consistently inhibits invasive and migratory behaviors, while AKT1 depletion reduces cellular proliferation and tumor growth (Figure 2D, Supplementary Figure 2O-T). We postulate that the AKT2-dependence of invasion and migration in these cells is regulated through the AKT2-specific transcriptional changes in EMT-related genes E-cadherin, ZEB1, and Snail (Figure 7). These data are consistent with findings in breast cancer, in which AKT2 but not AKT1 promotes cell migration and EMT [21,22,47,48], and the well-recognized role of EMT in metastatic promotion [49–51].

While a consistent and specific role for AKT3 was not identified in our study, this may be partially explained by our AKT3-specific knockdown efficiency, which varied greatly among cell lines (Supplementary Figure S3C). AKT3 amplification is known to occur in melanoma [26] but was not characterized across our panel, and could attenuate knockdown efficiency if present. AKT3 phosphorylation also appeared proportionally greater across cell lines (Supplementary Figure S2A), in partial agreement with previous reports that it is preferentially phosphorylated in melanoma [24]. Nevertheless, our finding that AKT2 knockdown greatly attenuates the metastatic potential of human melanoma cell lines is consistent with previous data suggesting that PHLPP1 inhibits melanoma metastasis by suppressing the phosphorylation of AKT2 and AKT3, but not AKT1 [25].

Once cells become migratory and invasive, extravasation and anchorage independent growth are two necessary requirements for metastatic colonization [49]. Previous work in PTEN-deficient prostate tumors demonstrated their dependence on AKT2 for both maintenance and survival, but AKT1 in the same tumors was dispensable [52]. Our xenograft studies suggest that PTEN-deficient melanomas are initially sensitive to AKT2 inhibition but ultimately do not depend on AKT2 long term given our observation of delayed tumor outgrowth in the AKT2 knock-down mice, as well as unmitigated tumor growth of AKT2 KO cells (Supplemental Figure S4B). In PTEN null melanomas and in the human cell lines in our study, a key difference is significant AKT1 phosphorylation, which could compensate to drive tumor cell growth and survival, given the cellular sensitivity to AKT1 depletion on cell proliferation and tumor growth that we observed (Supplementary Figure 6). Despite the apparent inability of AKT1 to fully support metastasis in AKT2-depleted human melanoma cells, hyperactive AKT1 contributes to the metastatic potential of the mouse SM1-750 model. Kircher and colleagues showed, utilizing an autochthonous mouse model of melanoma, that hyperactive, ectopic AKT1(E17K) resulted in enhanced level of brain metastases and reduced overall survival compared with hyperactivating mutations in AKT2 or AKT3, mediated through FAK, an AKT1-specific substrate [32]. Indeed, we observed a significant increase in the phosphorylation of both AKT1 and AKT3 after passage *in vivo* to generate the brain metastatic-competent SM1-750 cell line (Figure 1D), in addition to increased AKT2 phosphorylation (Figure 1F) despite the presence of intact PTEN protein (Supplementary FigureS1A). Whether this increased phosphorylation is due to hyperactivating mutations arising in AKT1 remains to be determined. Interestingly, Kircher and colleagues also show that mice with hyperactive AKT2 but not hyperactive AKT3 mutations still developed brain metastases, albeit to a lesser extent than AKT1. These observations, taken together with our data, suggest that AKT2 promotes melanoma metastasis in murine model systems, and further studies may elucidate mechanistic differences in metastatic promotion between AKT isoforms in murine melanomas.

To understand the potential stages of metastatic dissemination for which AKT2 may be most critical, we explored extravasation from blood vessels and tumor cell proliferation at seeded distant sites as two candidate stages for the study of metastasis. Data presented in this report provided strong evidence that the key limiting event regulated by AKT2 was tumor cell extravasation. By pre-incubating cells with doxycycline to knockdown AKT2 and then injecting them into the tail vein of mice, we observed complete lack of metastatic dissemination, despite the presence of phosphorylated AKT1 and AKT3. We ruled out the possibility that resident cells were dormant at distant metastatic sites by removing mice from doxycycline chow at 6 weeks. No tumors appeared after an additional 6-week observation period, suggesting disseminated cells may have been removed from circulation. It is also possible that 6 weeks was not sufficient time, or dormant cells were too small to observe by our methods, as melanoma cells disseminating to the brain have been known to exist as single cells or small clusters barely visible by histology that can then re-grow when conditions are optimal [53]. Nevertheless, when cells expressing AKT2 were injected into the tail vein and then mice were fed doxycycline chow to knock down AKT2 24h later, there was a partial inhibition of metastasis. However, these mice were not fully protected from metastatic disease, unlike the mice in inoculated with AKT2-depleted cells, suggesting AKT2 activity may be most impactful at the stage of extravasation, and its effect on tumor growth at the metastatic site was moderate. These results were phenocopied with AKT2 KO cells, in which AKT2 KO provided a survival benefit, perhaps by partially inhibiting the onset of metastatic disease.

We further investigated the mechanisms whereby AKT2 may be mediating these effects, hypothesizing that AKT2 KD cells might be glycolytically impaired. AKT2-specific roles in maintaining glucose homeostasis are well-documented [18], and metabolic re-wiring in melanoma can facilitate metastatic dissemination [54]. We observed that AKT2 KD suppressed basal glycolytic metabolism and reduced compensatory glycolysis. In malignant glioma, AKT2-specific phosphorylation of PDHK1 at Thr346 was shown to increase phosphorylation of PDHE1α. This interferes with the entry of pyruvate into the TCA cycle, resulting in stimulation of glycolysis, the maintenance of cell proliferation, and the inhibition of autophagy and apoptosis during severe hypoxia [42]. Earlier studies have shown that AKT2 selectively promotes the expression of miR-21 during hypoxia and renders cells resistant to hypoxia-induced cell death [55] Importantly, recent studies have shown that miR-21 promotes glycolysis by targeting pyruvate dehydrogenase A1 (PDHA1) [56] Therefore, AKT2 may inhibit the activity of pyruvate dehydrogenase by increasing the expression and/or activity of PDHK1 and inhibiting the expression of PDHA1. In agreement with a significant role for AKT2-mediated regulation of PDHK1 and subsequent PDHE1α activity, we observed reductions in PDHK1 as well as phosphorylated PDHE1α with AKT2 KD. Taken together, the totality of our data suggests AKT2-mediated regulation of PDHK1 and other components of pyruvate metabolism are important in both normal and hypoxic conditions, although the specific regulation may differ. Future studies should further define a role for PDHK1 in glycolytic maintenance mediating melanoma metastasis.

## 5. Conclusion

In summary, using genetically engineered human cell lines and novel syngeneic mouse models, we reveal multiple roles for AKT2 in melanoma metastasis. Indeed, as AKTs are central pleiotropic signaling hubs [57], it is not surprising that AKT2-mediated cellular changes facilitating melanoma metastasis are manifold, including glycolytic changes and enhancement of factors promoting the epithelial to mesenchymal transition. This study reinforces the need for improved development of clinically relevant small molecules for selective targeting of AKT isoforms as melanoma therapy.

## Author Contributions

S.M designed and performed experiments, analyzed data, and drafted the manuscript. J.P performed experiments and edited the manuscript. A.L.B performed experiments, and drafted the manuscript. P.T. assisted with experimental design and interpretation of results. P.W.H. initiated the project, designed experiments, and helped to write and edit the manuscript.

## Data Availability Statement

Data available within the article or on request from the authors

## Acknowledgements

We gratefully acknowledge all collaborators that provided reagents and materials (as indicated in materials and methods). We thank Drs. Gary Sahagian, Min Fang of the Small Animal Imaging Facility, and Alessandra Cecchini for help with *in vivo* imaging. Model figures made with Biorendr.

