## Supplementary material for "AKT2 Loss Impairs BRAF-Mutant Melanoma Metastasis": Tables S1-S2 and Figures S1-S6

### Supplementary Materials:

**Listed:**

Table S1: Primer Sequences and Hairpin Sequences

Table S2: Antibodies Used

Figure S1:. Characterization of AKT Phosphorylation and Knockdown Cell Lines

Figure S2: AKT2 knockdown impairs wound healing, migration and invasion in WM455 and UACC903 cells.

Figure S3: Activation state of AKT isoforms in murine melanoma cells *in vitro* and *in vivo*.

Figure S4: AKT1 Deletion Delays Primary Melanoma Growth and Improves Survival

Figure S5: AKT1 Knockdown impairs Melanoma Cell Proliferation

Figure S6: AKT1 Knockdown Restricts Anchorage Independent Growth

| qPCR Target | Forward | Reverse |
|---|---|---|
| ECAD | GAACGCATTGCCACATACAC | GAATTCGGGCTTGTTGTCAT |
| ZEB1 | GCACCTGAAGAGGACCAGAG | TGCATCTGGTGTTCCATTTT |
| MMP2 | CCGTCGCCCATCATCAAGTT | CTGTCTGGGGCAGTCCAAAG |
| Snail | CACTATGCCGCGCTCTTT | GGTCGTAGGGCTGCTGGAA |
| TEAD2 | CTCACTCCGTAGAAGCCACC | TGCCTTCTTCCTGGTCAAGT |
| TEAD3 | GCACCTTCTTCCGAGCTAGA | TACGGCCGAATGAGTTGATT |
| TBP | GAGCCAAGAGTGAAGAACAGTC | GCTCCCCACCATATTCTGAATCT |
| **shRNA Hairpins** | | |
| AKT1 | CCGGGAGTTTGAGTACCTGAAGCTGCTCGAGCAGCTTCAGGTACTCAAACTCTTTTTG | |
| AKT2 | CCGGGCGTGGTGAATACATCAAGACCTCGAGGTCTTGATGTATTCACCACGCTTTTTG | |
| AKT3 | CCGGCTGCCTTGGACTATCTACATTCTCGAGAATGTAGATAGTCCAAGGCAGTTTTTG | |
| Non-Targeting | CCGGCAACAAGATGAAGAGCACCAACTCGAGTTGGTGCTCTTCATCTTGTTGTTTTT | |

**Table S1: Primer Sequences and Hairpin Sequences**

| Antibody | Source | Catalog Number |
|---|---|---|
| AKT1 | Cell Signaling Technologies | 2938 |
| AKT2 | Cell Signaling Technologies | 5239 |
| AKT3 | Cell Signaling Technologies | 8018 |
| Alpha Tubulin | Cell Signaling Technologies | 3873 |
| Beta Actin | Sigma | A5441 |
| p-RXRXXS/T | Cell Signaling Technologies | 9611 |
| pan AKT | Cell Signaling Technologies | 4691 |
| pan phospho-AKT | Cell Signaling Technologies | 4060 |
| PDHE1a | Cell Signaling Technologies | 3205 |
| PDHK1 | Cell Signaling Technologies | 3820 |
| phospho-AKT1 | Cell Signaling Technologies | 9018 |
| phospho-AKT2 | Cell Signaling Technologies | 8599 |
| phospho-PDHE1a | Cell Signaling Technologies | 31866 |
| PTEN | Cell Signaling Technologies | 9552 |

**Table S2:** Antibodies Used

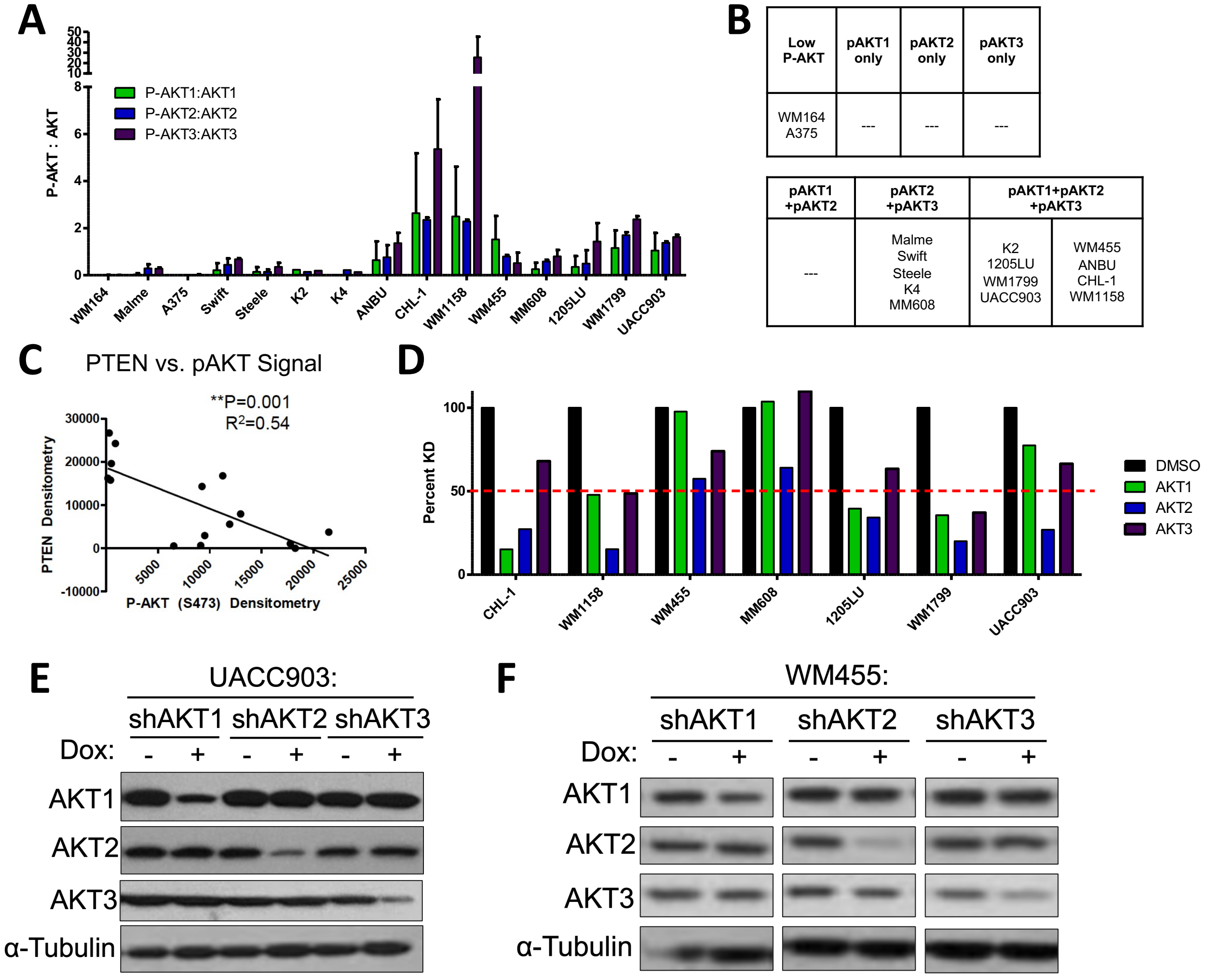

**Supplementary Figure 1. Characterization of AKT Phosphorylation and Knockdown Cell Lines**

**A.** Quantitation of immunoblotting by ImageJ for AKT isoform-specific phosphorylation normalized to phospho-protein over total. **B.** Summary of isoform specific AKT phosphorylation across human cell lines. **C.** Analysis of densitometry of immunoblotting comparing total AKT phosphorylation (S473) to PTEN protein level. **D.** Akt-isoform knockdown efficiency was quantified from western blots using ImageJ in novel generated human melanoma cell lines after 72h of DOX treatment (0.5ug/mL) and normalized to DMSO-treated and loading controls. **E-F.** Representative immunoblots of human melanoma cell lines UACC903 **(E)** and WM455 **(F)** showing Akt-isoform knockdown efficiency after 72h DMSO or doxycycline exposure (0.5ug/mL).

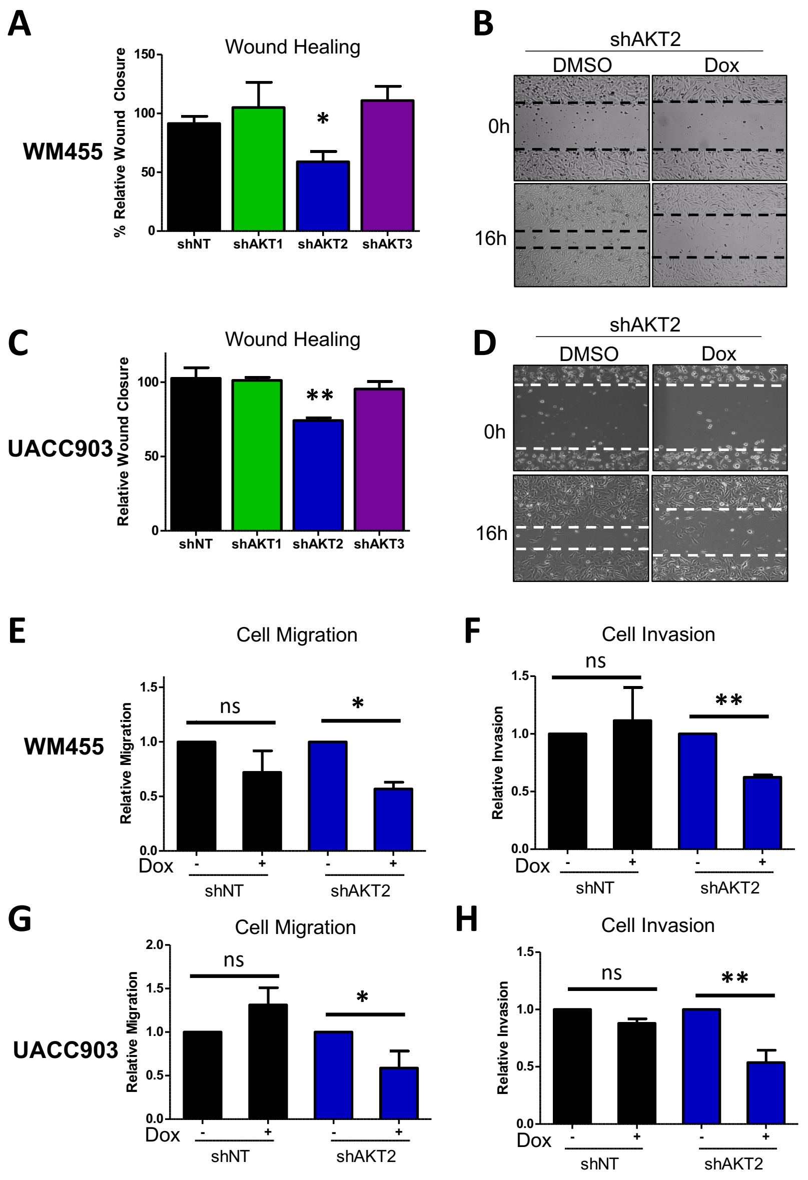

**Supplementary Figure 2. AKT2 knockdown impairs wound healing, migration and invasion in WM455 and UACC903 cells.**

**A-D.** Wound closure in indicated cell lines at 0h and 16h post scratch in AKT2 KD cells treated with DMSO- or doxycycline (DOX)-containing media. Shown are quantification for DOX relative to DMSO treated WM455 cells (**A**) or UACC903 cells (**C**), with representative images of control (NT) or isoform-specific KD cells treated with DOX or DMSO at 0 and 16 hours for WM455 cells (**B**) or UACC903 cells. Cell migration ability after treatment in DMSO or DOX containing media using transwell assay for WM455 cells (**E**) or UACC903 cells (**G**). Cell invasion through Matrigel coated membranes after treatment in DMSO or DOX containing media for WM455 cells (**F**) or UACC903 cells (**H**).

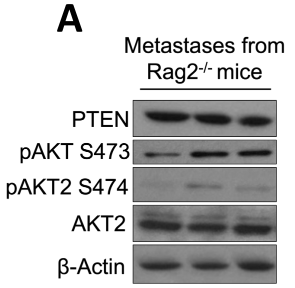

B

| Cell Line | BRAF$^{V600E}$ | INK4A | ARF | Notes |
|---|---|---|---|---|
| SM1 | + | WT | WT | Generous gift from Toni Ribas |
| SM1-750 | + | WT | WT | SM1 cells passaged in BL6 mice and infected with pLENTI-luciferase |
| TUMM4218 | + | -/- | -/- | |
| TUMM4228 | + | WT | -/- | |
| TUMM4246 | + | WT | -/- | |
| TUMM8253 | + | WT | -/- | Mouse received UV radiation at PND 3.5 |

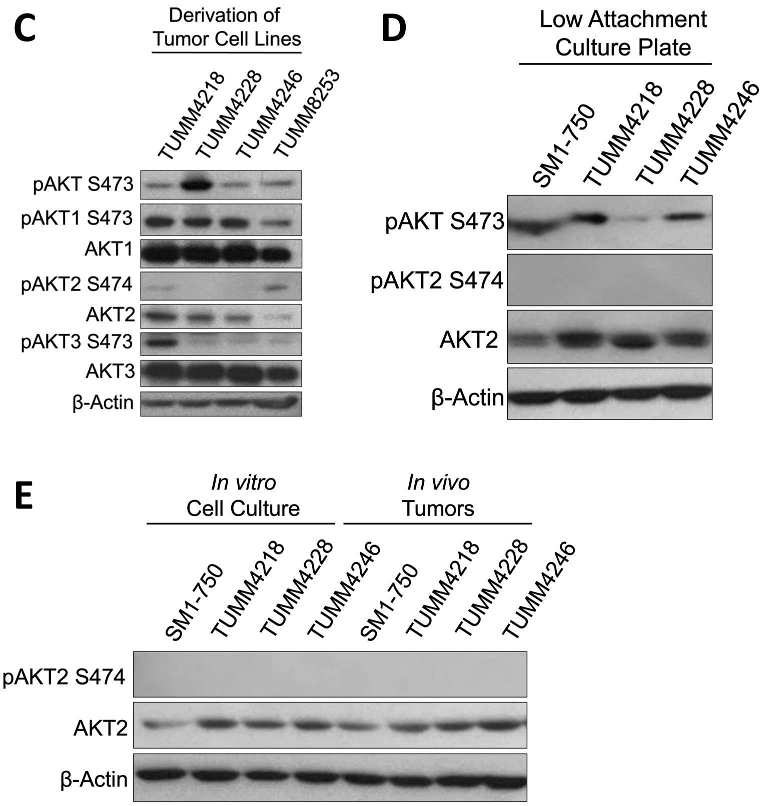

**Supplementary Figure 3. Activation state of AKT isoforms in murine melanoma cells *in vitro* and *in vivo*. A.** SM1-750 cells were injected into the tail vein of Rag2-/- mice and metastatic nodules were collected after 4 weeks, or when mice became moribund, homogenized and subjected to immunoblotting. **B.** Table characterizing SM1 and SM1-750 derived cells (Figure 3) and lines derived form spontaneous primary melanomas in mice expressing human BRAFV600E transgene after isolation and passage in culture. TUMM: Tufts University Mouse Melanoma. PND: post-natal day **C.** Expression and phosphorylation of each AKT isoform was characterized by immunoblotting. **D.** SM1-750 and TUMM cell lines were cultured in low-attachment plates until macroscopic colonies were visible, then collected and subjected to immunoblotting. **E.** SM1-750 and TUMM cell lines were grown in standard 2-D adherent culture, or injected subcutaneously into NODSCID mice and the resulting tumors collected and subjected to immunoblotting with indicated antibodies.

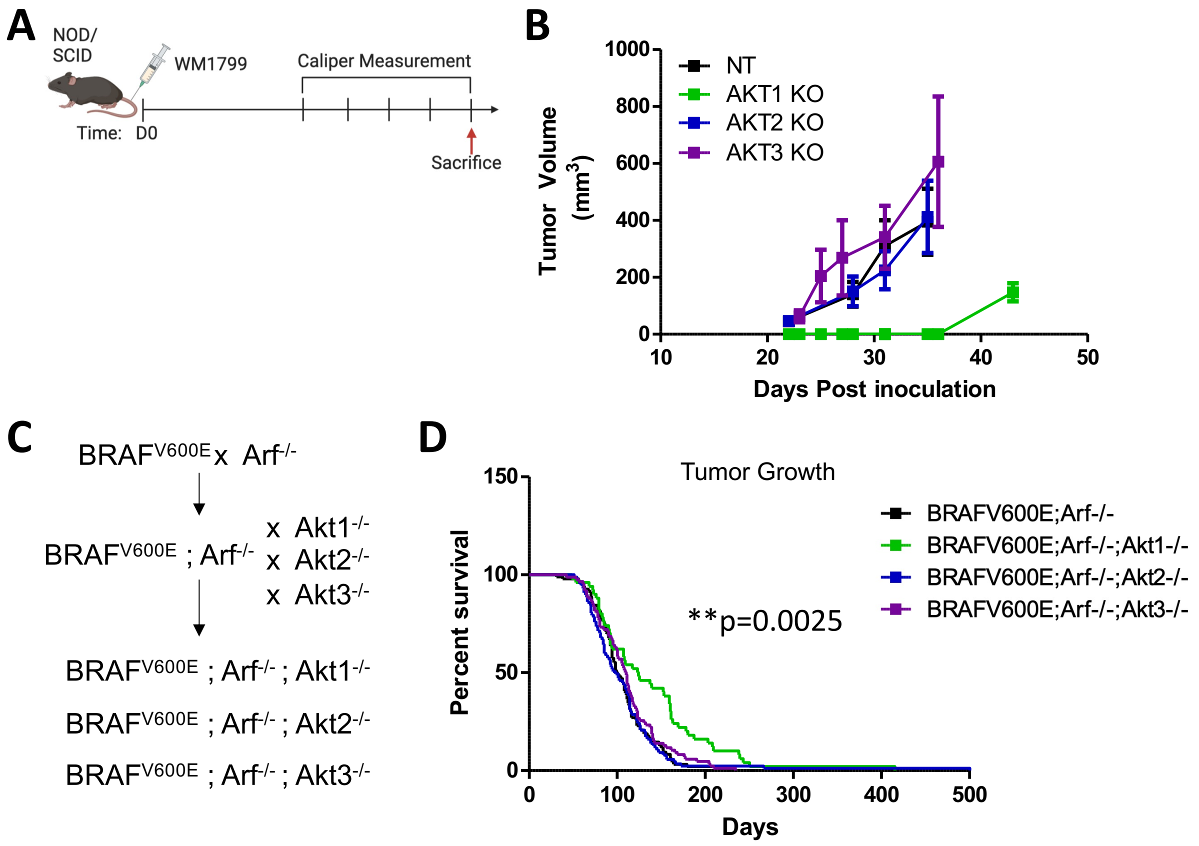

**Supplementary Figure 4: AKT1 Deletion Delays Primary Melanoma Growth and Improves Survival.** A. Experimental schematic in which NT or AKT isoform specific WM1799 cells were injected into the flank of NOD/SCID mice and tumor growth was monitored by Caliper measurement over time, quantified in B. C. Breeding scheme for creation of melanoma prone Akt isoform KO mice by cross with $BRAF^{V600E}ARF^{-}$/- mice. D. Survival curve representing percent overall survival of AKT isoform KO melanoma prone mice over time.

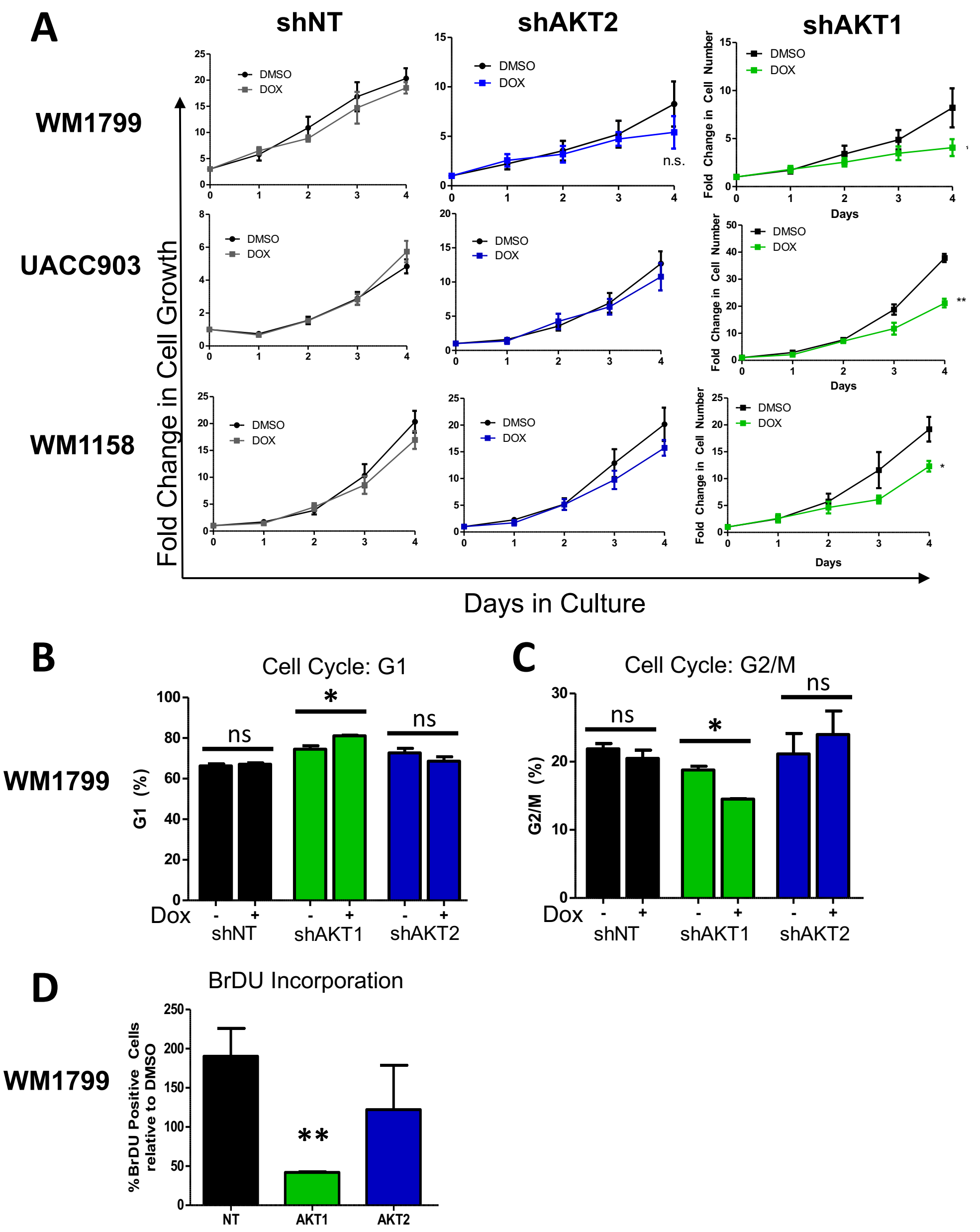

**Supplementary Figure 5. AKT1 Knockdown impairs Melanoma Cell Proliferation. A.** Cell proliferation of human melanoma cell lines expressing non-targeting (shNT), AKT2 (shAKT2), or AKT1 (shAKT1) hairpins in the presence of DMSO- or DOX-containing media assessed using cell counting with trypan blue exclusion and represented as a fold change from day 0 over 4 days. **B-C.** Cell cycle analysis by propidium iodide staining to assess G1 or G2/M fraction of WM1799 cells expressing shNT or shAKT isoform hairpins in the presence (+) or absence (-) of DOX-containing media. **D.** BrDu incorporation was quantified from WM1799 cells expressing inducible AKT-isoform or NT hairpins and grown in the presence of DOX or DMSO for 2 days before adding BrDu for one hour, plotted relative to DMSO treated cells.

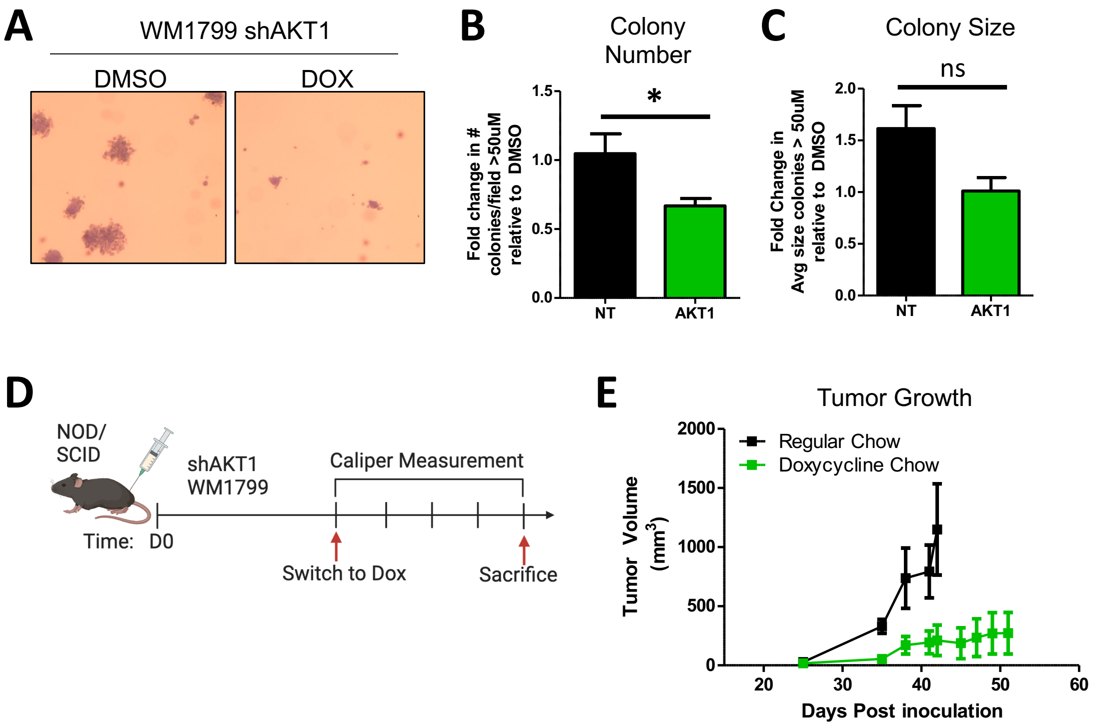

**Supplementary Figure 6. AKT1 Knockdown Restricts Anchorage Independent Growth A.** Anchorage-independent growth of WM1799 shAKT1 cells in soft-agar and incubated with DMSO or DOX-containing media. Colonies were fixed and stained with crystal violet and colony number **(B)** or colony size **(C)** greater than 50uM were counted using ImageJ. **D.** Experimental scheme showing WM1799 shAKT1 cells were injected subcutaneously into NODSCID mice and maintained on either regular or DOX containing chow. Mice were sacrificed when tumors reached 1500m$^3$ according to approved protocols. **E.** Tumor growth by caliper measurement was quantified over time.
